# AchE-OGT dual inhibitors: Potential Partners in Handling Alzheimer’s Disease

**DOI:** 10.1101/303040

**Authors:** Mukta Sharma, Ashwin K. Jainarayanan

## Abstract

The emerging role of O-GlcNAc Transferase (OGT) and tau protein in Alzheimer’s disease (AD) holds promises for the treatment of this life-threatening neurodegenerative disorder. In view of the availability of 3D structure of OGT, we attempted to develop structure-based pharmacophore model to elucidate specific structural requirements for binding of inhibitors to the active site of OGT. During the course of study we discovered that donepezil, an old and trusted Acetylcholinesterase (AchE) inhibitor also possess the pharmacophoric sites important for interaction with OGT active site. To further explore the specific interactions of donepezil with OGT, we performed molecular docking studies using CDocker. The results of molecular docking and structure-based pharmacophore mapping revealed that donepezil has the required structural features to interact with various OGT active site amino acids like Lys842, His920, Leu653, Gly654, Asn838, and Thr921 in addition to its AchE interaction abilities. Our findings could be an important breakthrough in the design of OGT specific and/or AchE OGT dual inhibitors.

## Introduction

Donepezil has long been recognized as a potent reversible inhibitor of enzyme AchE [1, 2] making it a useful drug in the treatment of Alzheimer’s disease (AD). The complexed structure of donepezil and AchE is freely available through Protein Data Bank site (PDB ID: 3PE4) [3]. On the other hand post-translational modification of serines/threonines [4] containing proteins by transfer of N-acetyl glucosamine have gained attention for their prospective role in neurodegenerative diseases like the AD. Essentially it is now known that OGT is responsible for this enzymatic reaction and moreover now there is sufficient evidence to link OGT and tau [5] protein. The crystal structures of human OGT [3] containing 4.5 tetratricopeptide repeat (TPR) units and the catalytic domain (hOGT4.5) has been resolved. 3PE3 is the complex of OGT and UDP resolved at 2.8 Å, whereas 3PE4 is a complex containing UDP and a well-characterized acceptor, 14-residue CKII peptide substrate, resolved at 1.95 Å. Both the structures are accessible through PDB site. The emerging role of OGT [6] in AD and availability of its 3D structure prompted us to undertake detailed structure-based pharmacophore study for better understanding of OGT binding site which would help in the design of OGT inhibitor/substrate.

## Methods

### Structure-based pharmacophore modeling

The three-dimensional structure of OGT (3PE4) complexed with UDP was obtained from PDB database and was used to develop pharmacophore model to uncover the putative binding site and structural requirement of the OGT inhibitors.

The protein structure was monitored [7] for valence and the missing hydrogens were added, the structure was further checked using protein health check tool for any structural error. The cleaned enzyme structure was subjected to active site identification. The receptor’s active site was identified using a sphere whose location and radius was adjusted to 9.0 Å, so as to include the active site and the key residues of the protein involved in interaction with ligands.

Keeping the density of lipophilic sites and density of polar sites parameter value to 10, the interaction map was generated. The interaction map often displays a large number of features, especially when the protein is capable of binding a variety of ligands and has a number of different binding modes. Thus, deriving pharmacophore models directly from the interaction map can be quite complicated. To overcome this problem, neighboring features of the same type were grouped into the same cluster. The feature which was closest to the geometric center of the cluster was selected to represent the cluster, whereas the rest of the features were omitted. However, even after clustering the numbers of the features were still too high to use all of them in a single query. A query composed of all the features may fail to retrieve any hits from the database/compound library. Therefore, multiple 3D queries, composed of fewer numbers of features, were generated from the interaction map by considering all the possible combinations. The final model constructed was subjected to non-feature atoms exclusion. The exclusion constraint feature is an object that represents an excluded volume in space, within a given radius. The excluded volumes were placed on regions of space that are occupied by the inactive molecules but not the active molecules. A pharmacophore with an excluded volume only matches if no atoms penetrate the excluded area. The final hypothesis contained six features: two hydrogen bond donors, two hydrogen bond acceptors and two hydrophobic groups describing the interactions between the protein OGT and UDP.

Accuracy and predictability of the developed model was evaluated by mapping some of the known OGT substrates/inhibitors onto the developed pharmacophore. The compounds which mapped very well onto the developed structure-based pharmacophore included thiamet-G [8] a known inhibitor of OGT and UDP the usual substrate for OGT. During our study, we found that while mapping donepezil onto the OGT driven pharmacophore, it showed a four-feature mapping. Since this was an unobvious and surprising outcome, our team decided to further explore the interaction of donepezil with OGT through docking studies.

### Molecular Docking

The structure-based docking of donepezil to the active site of OGT was carried out using the CDocker [9] program, which is a molecular dynamics simulated annealing based algorithm [10].

The crystal structure of OGT comprising of 4 chains (A, B, C and D) was obtained from PDB and checked for valency, missing hydrogens and any structural disorders like connectivity and bond orders. Water molecules from the protein hierarchy were removed and only A chain with its crystal ligand was retained as this chain has the active site center. The A chain was split into the protein and ligand part. The protein was defined as receptor molecule whereas crystal ligand was used to define the binding site of 9 Angstroms on the receptor molecule. The structure of test compound donepezil was sketched and energy minimized to obtain most stable structure to be used for docking. Based on UDP coordinates donepezil was docked into the active site of OGT with all the parameters set to their default values. Finally, all the possible interaction modes for different alignments were analyzed on the basis of CDOCKER interaction energies.

## Results and Discussion

Twenty poses were obtained for donepezil with the CDocker interaction energy ranging between 33 and 47. All the poses were inspected for the type of interaction between donepezil and binding site of OGT. It was observed that one of the methoxy group attached to the indene ring of donepezil (Figure 1) is interacting with Lys842, His920, and Lys430 (Figure 2(a-c)). In line to this, structure-based pharmacophore mapping results also showed hydrogen bonding interaction with methoxy group (Figure 3; green color coded for hydrogen bond acceptor feature). The other methoxy group also showed a close hydrogen bonding interaction with Lys430 and His920 (Figure 2(b, c)) whereas, structure-based pharmacophore revealed that this methoxy group is involved in hydrophobic interactions (Figure 3; cyan color coded for hydrophobic feature). Though at this point the results of structure-based pharmacophore and docking were not in agreement, a close inspection of nearby amino acids confirmed the presence of hydrophobic amino acids like Phe868 and Leu1009. Hence, the presence of hydrogen bonding and hydrophobic amino acids close to second methoxy group clearly points towards hydrophobic and hydrogen bonding interaction of this group with the active site of OGT. As shown by structure-based pharmacophore mapping the carbonyl oxygen of indene ring is involved in hydrogen bonding (Figure 3; green color coded for hydrogen bond acceptor feature).

**Figure 1.**
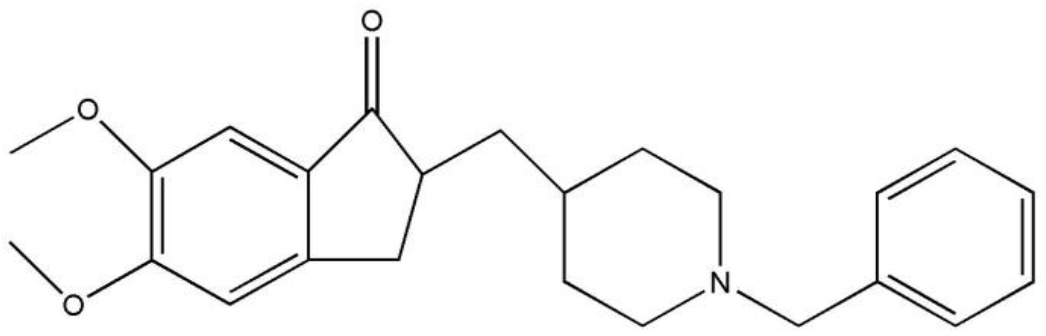
Structure of donepezil.

**Figure 2.**
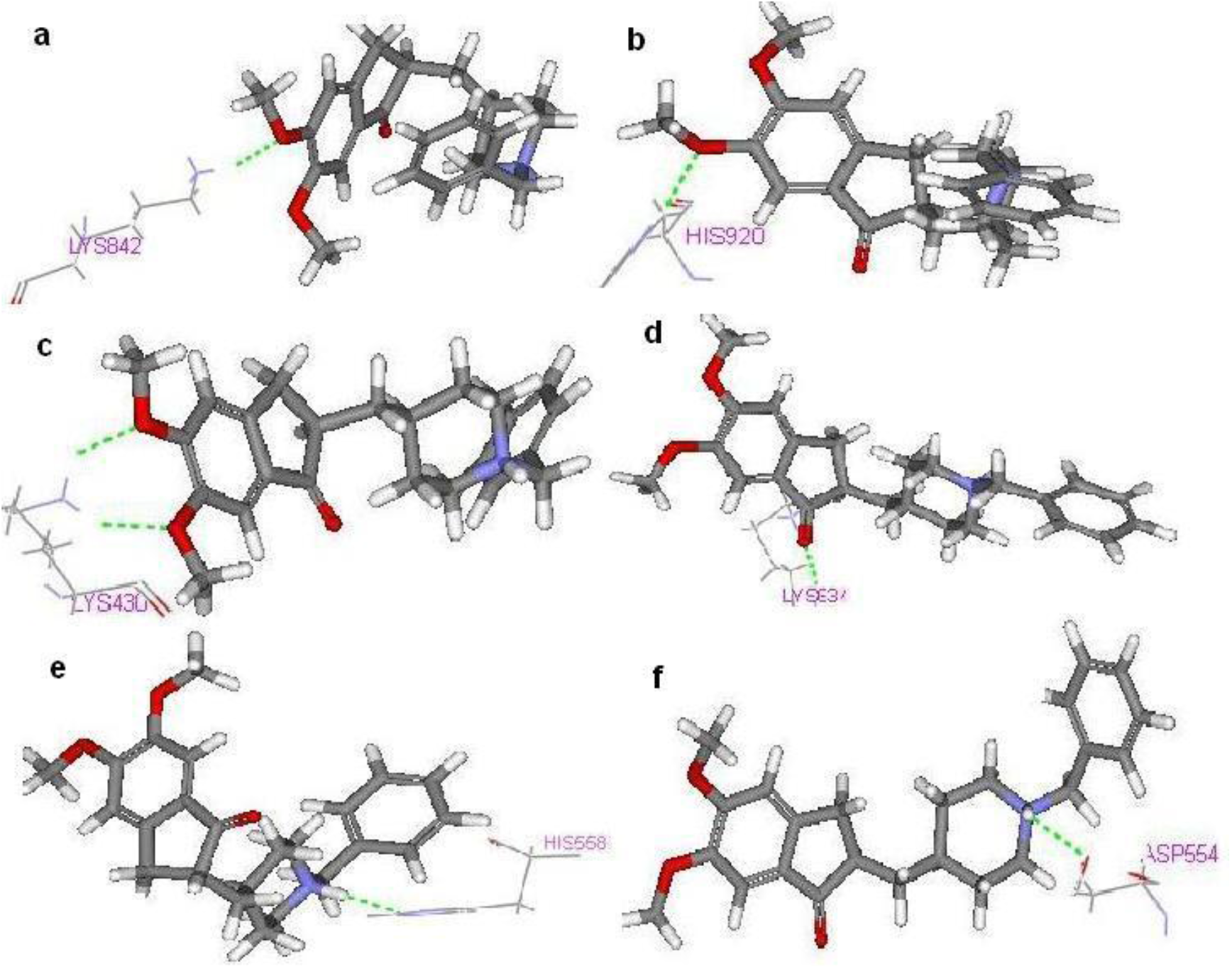
Docked conformation of donepezil on to the active sites of OGT. Dotted lines represent Hydrogen bonds (green color). [(**a**) Interaction of Lys842 with methoxy group attached to 6^th^ position of indene ring of donepezil (**b**) Interaction of His920 with methoxy group attached to 6^th^ position of indene ring of donepezil (**c**) Interaction of Lys430 with methoxy group at 5^th^ and 6^th^ positions of donepezil (**d**) Interaction of Lys634 with carbonyl oxygen of donepezil (**e**) Interaction of Asp554 with NH of benzene ring of donepezil (**f**) Interaction of His558 with NH of benzene ring of donepezil].

**Figure 3.**
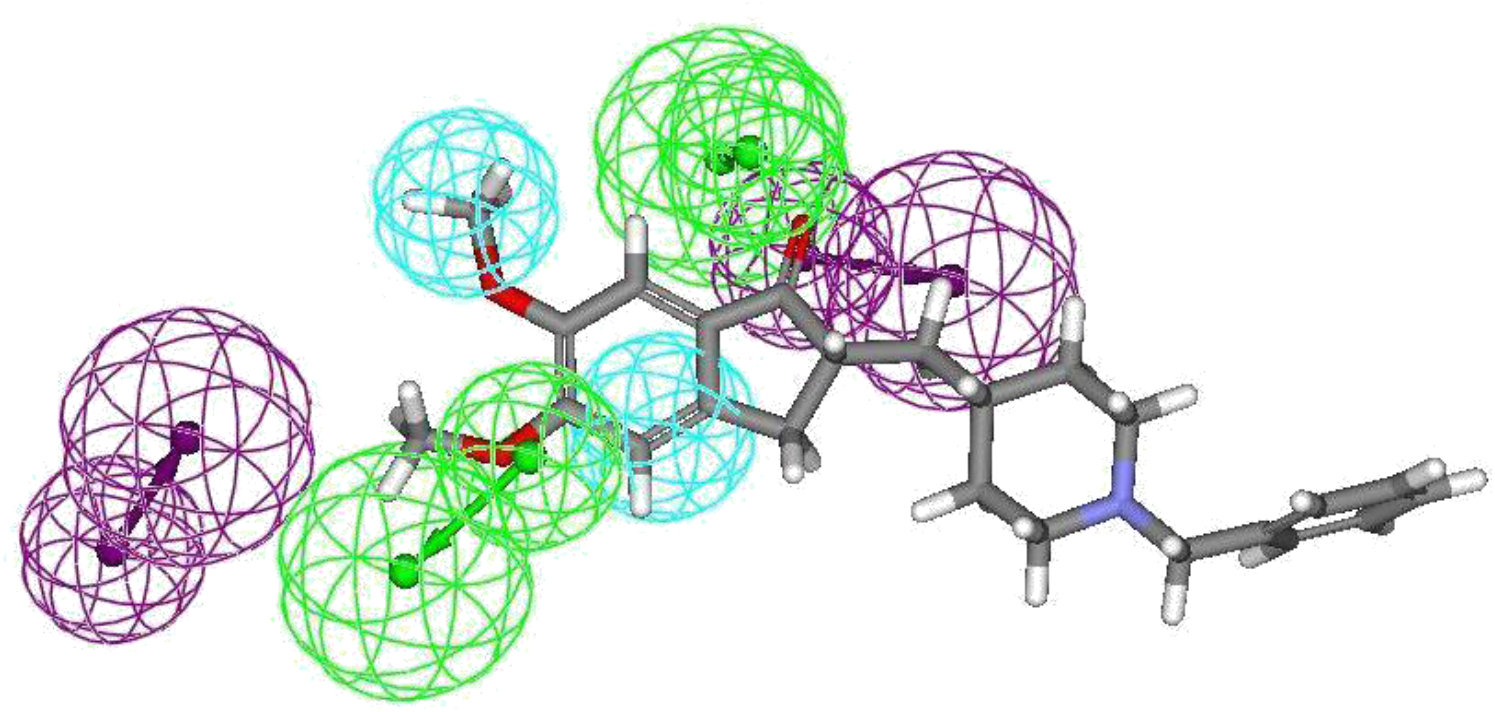
Four feature pharmacophore mapping of donepezil on OGT active site driven pharmacophore. The mapped features are color-coded with cyan, hydrophobic; green, hydrogen bond acceptor; magenta, hydrogen bond donor.

This was further in agreement to docking results which the showed interaction of carbonyl oxygen with Lys634 (Figure 2(d)). The second hydrophobic interaction appearing in structure-based pharmacophore (Figure 3; cyan color coded for hydrophobic interactions) clearly showed the involvement of terminal phenyl ring and this was also confirmed by the presence of hydrophobic amino acids like Leu653, Phe693 during docking. While analyzing the docking results, it was also observed that NH of the benzene ring is involved in hydrogen bonding with His558 and Asp554 (Figure 2(e, f)). These results clearly suggest that donepezil structure has required pharmacophoric sites to interact with OGT.

Review of previous literature of donepezil and AchE interaction revealed that the terminal phenyl ring of donepezil interacts with Trp86A, the indene ring of donepezil interacts with hydrophobic amino acids including Trp286A, and π-π interactions have been observed between terminal phenyl ring and indene ring of donepezil and Trp286A. In addition to this, there are other hydrophobic interactions between Tyr337A, Trp286A, and Phe338A, Tyr341A, Trp86A, and functional groups of donepezil. In fact on comparison of donepezil-OGT interaction results with previously reported donepezil-AchE interaction [11] data, it was observed that in both the cases the pharmacophoric sites and nature of interaction are different indicating the possibility of dual interaction ability of donepezil. This was further established by thorough analysis of the inter-feature distances (Angstrom) obtained from our in-house developed structure based pharmacophore of OGT and AchE. The results revealed that in both the cases the interfeature distances are different (Figure 4 and Figure 5) but it is the uniqueness of the donepezil molecule in terms of essential features at required distances, which ensures its interaction ability with AchE and OGT.

**Figure 4.**
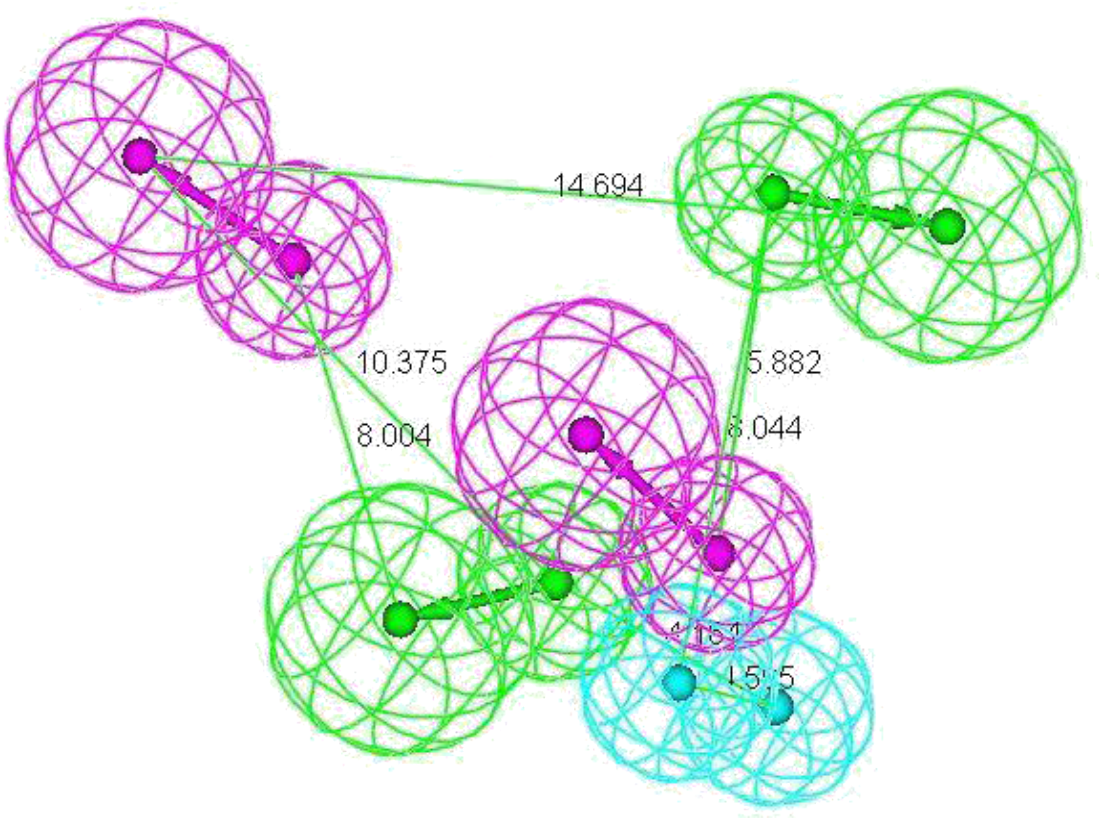
The AchE pharmacophore with inter-feature distances in Angstrom. The features are color coded with cyan, hydrophobic; green, hydrogen bond acceptor; magenta, hydrogen bond donor.

**Figure 5.**
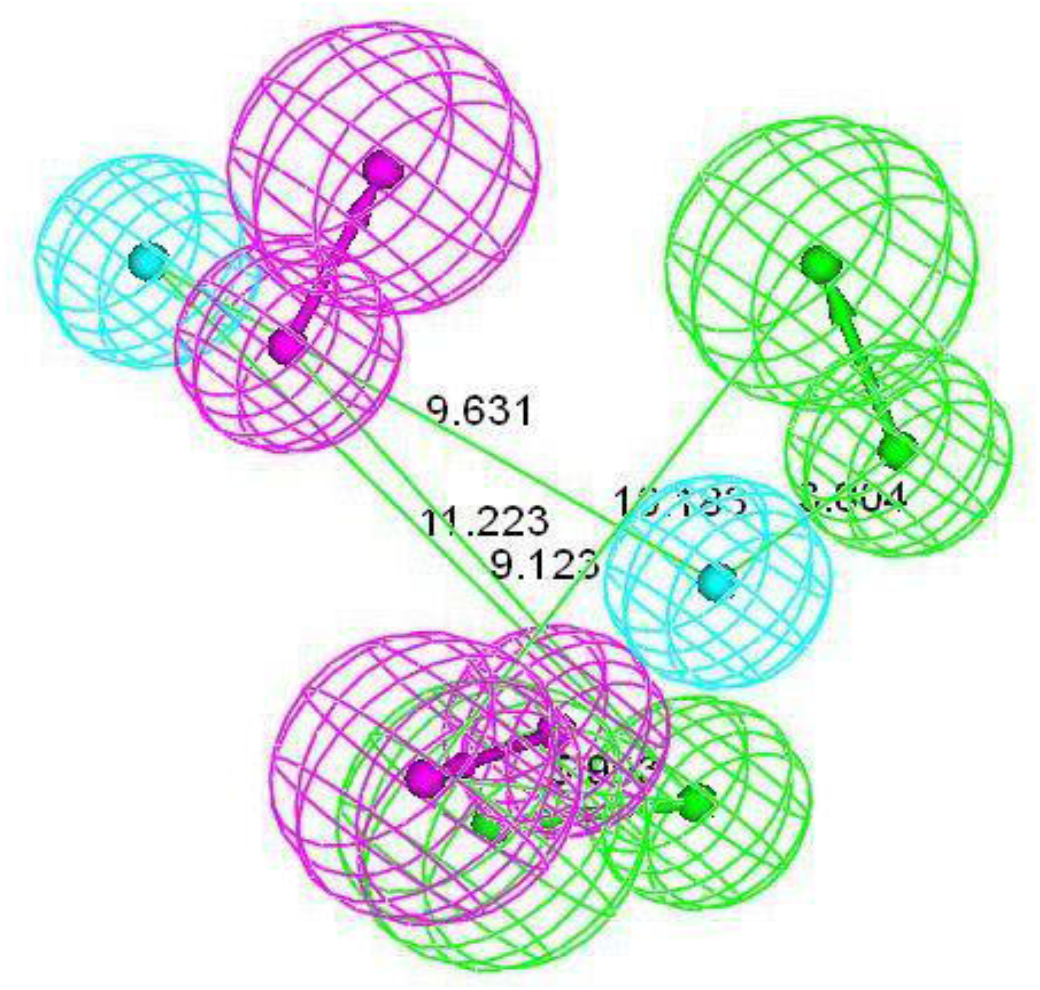
The chemical features of OGT pharmacophore with their inter-feature distances in Angstrom. The features are color-coded with cyan, hydrophobic; green, hydrogen bond acceptor; magenta, hydrogen bond donor.

## Conclusions

The findings of structure-based pharmacophore mapping and molecular docking clearly show that donepezil has the required structural features to interact with various OGT active site amino acids like Lys842, His920, Leu653, Gly654, Asn838, and Thr921 in addition to its AchE interaction abilities. In view of this, it can be concluded that donepezil an old and trusted AchE inhibitor has pharmacologically related and significant off-target OGT interaction capabilities. This study could serve as an ideal guide to scientific community focusing on design of potent and specific OGT and/or prospective AchE-OGT dual inhibitors.

## Acknowledgement

We are grateful to Prof. Aditya Shastri, Vice Chancellor, Banasthali University (India) for providing necessary computational facilities for the study.

## Author contributions

MS performed the whole simulation experiment and had made major contribution to conception and design of the experiments. AKJ supported in analyzing the results and composing the manuscript. MS and AKJ contributed to draft the manuscript. Both authors have read and approved the final manuscript.

## References

[1] Z. Fu, X. Li, Y. Miao, and K.M. Merz Jr., Conformational analysis and parallel QM/MM X-ray refinement of protein bound anti-Alzheimer drug donepezil, J. Chem. Theory Comput. 9 (2013), pp. 1686–1693.

[2] G. Kryger, I. Silman, and J.L. Sussman, Structure of acetylcholinesterase complexed with E2020 (Aricept): implications for the design of new anti-Alzheimer drugs, Structure 7 (1999), pp. 297307.

[3] Y. Nam, J. Jiang, P. Sliz, and S. Walker, Structure of human O-GlcNAc transferase and its complex with a peptide substrate, Nature 469 (2011), pp. 564–567.

[4] L. Wells, L.K. Kreppel, F.I. Comer, B.E. Wadzinski, and G.W. Hart, O-GlcNAc transferase is in a functional complex with protein phosphatase 1 catalytic subunits, J. Biol. Chem. 279 (2004), pp. 38466–38470.

[5] L. Buée, T. Bussière, V. Buée-Scherrer, A. Delacourte, and P.R. Hof, Tau protein isoforms, phosphorylation and role in neurodegenerative disorders, Brain Res. Rev. 33 (2000), pp. 95–130.

[6] L.K. Kreppel, M.A. Blomberg, and G.W. Hart, Dynamic glycosylation of nuclear and cytosolic proteins, J. Biol. Chem. 272 (1997), pp. 9308–9315.

[7] S. Paliwal, D. Yadav, R. Yadav, and S. Paliwal, In silico structure-based drug design approach to develop novel pharmacophore model of human peroxisome proliferator - activated receptor γ agonists, Med. Chem. Res. 20 (2011), pp. 656–659.

[8] S.A. Yuzwa, M.S. Macauley, J.E. Heinonen, X. Shan, R.J. Dennis, Y.H. Garrett, E. Whitworth, K.A. Stubbs, E.J. McEachern, G.J. Davies, and D.J. Vocadlo, A potent mechanism-inspired O-GlcNAcase inhibitor that blocks phosphorylation of tau in vivo, Nat. Chem. Biol. 4 (2008), pp. 483–490.

[9] G.S. Wu, D.H. Robertson, C.L. Brooks, and M. Vieth. Detailed analysis of grid-based molecular docking: A case study of CDOCKER—A CHARMm-based MD docking algorithm, J. Comput. Chem. 24 (2003), pp. 1549–1562.

[10] Discovery Studio 2.1. Accelrys, Inc., San Diego, CA, 2005.

[11] J. Cheung, J.M. Rudolph, F. Burshteyn, S.M. Cassidy, N.E. Gary, J. Love, C.M. Franklin, and J.J. Height, Structures of human acetylcholinesterase in complex with pharmacologically important ligands, J. Med. Chem. 55 (2012), pp. 10282–10286.

